# Sonic hedgehog is required for neural crest-dependent patterning of the intrinsic tongue musculature

**DOI:** 10.1101/154443

**Authors:** Shigeru Okuhara, Anahid A. Birjandi, Hadeel Adel Al-Lami, Tomoko Sagai, Takanori Amano, Toshihiko Shiroishi, Karen J. Liu, Martyn T. Cobourne, Sachiko Iseki

## Abstract

The tongue is a highly specialized muscular organ important for breathing, speech, taste and swallowing. The secreted signaling molecule Sonic hedgehog (Shh) is expressed in dorsal tongue epithelium from the initial developmental stage. In this study, we utilized a series of genetic approaches to investigate the role of Shh during mouse tongue formation. Temporal-specific global deletion of *Shh* demonstrated a functional requirement for normal patterning of the intrinsic tongue muscles and establishment of the lingual tendon. These defects were reproduced in the mutant with a specific loss of signaling in oropharyngeal epithelium by a *Shh cis*-enhancer. In these mutants, *Ptch1* was lost in the underlying cranial neural crest (CNC)-derived mesenchymal lineage. The importance of Shh was confirmed by generating tissue-specific deletions in the ciliopathy gene *Ofd1*, which transduces Shh signaling. These results revealed Shh roles in patterning of the mesodermal intrinsic tongue muscles through CNC-derived mesenchyme, including the lingual tendon.

## Introduction

The tongue is a highly specialized muscular organ, which is required for normal mandibular growth and palatogenesis in the developing embryo and essential for airway maintenance, phonetic articulation, oral sensation and swallowing in post-natal life. Correct functioning of the tongue requires the cooperation of extrinsic and intrinsic muscles, tendons, neurons and a generous vasculature. Despite the functional importance of the tongue, comparatively little is known about the signaling events required to coordinate all these component structures to the development.

In the embryo, tongue primordia appear as paired swellings on the ventral wall of the pharyngeal arches at around embryonic day (E)10 in mice. These swellings grow and ultimately fuse with a medial lingual swelling to become a single midline structure in the floor of the early oral cavity. Myoblasts populate these early primordia following migration from the hypoglossal cord, a condensed region of mesoderm lying at the ventral edges of the occipital myotomes (Noden and West, 2006). The process of subsequent tongue development requires tissue interactions between the oropharyngeal epithelium, cranial neural crest (CNC)-derived mesenchyme and myogenic mesoderm (Parada et al., 2012).

Sonic hedgehog (Shh) is a secreted protein that plays a key role in diverse biological events extending from early development through to post-natal tissue homeostasis (Xavier et al., 2016; Peng and Joyner, 2015; Zuniga, 2015; Thesleff and Sharpe, 1997). Signal transduction is coordinated primarily through direct ligand-dependent inhibition of Patched-1 (Ptch1), a multi-span transmembrane protein that in the resting state indirectly inhibits a further multi-span protein Smoothened (Smo), which is absolutely required for transduction. These events are coordinated at the primary cilium and ultimately lead to pathway activation through post-transcriptional modification of Gli protein activity (Chang et al., 2016; Tabler et al., 2016). During embryonic development, *Shh* is expressed in very specific domains such as the node, notochord and floor plate, as well as in the limbs. Early expression of *Shh* in the midline is crucial for normal craniofacial development with complete loss of *Shh* function leading to holoprosencephaly (Gregory et al., 2015; Chiang et al., 1996) and a catastrophic failure of orofacial patterning. More recent studies have begun to distinguish the individual spatio-temporal signaling functions of *Shh* during craniofacial development (Xavier et al., 2016). Removal of Hedgehog (Hh)-responsiveness from CNC-derived mesenchyme through deletion of *Smo* in CNC cells (*Wnt1-Cre*; *Smo^flox/-^*) has demonstrated a requirement for Hh signaling in this population within the early facial processes from their commitment stage (Jeong et al., 2004). Significantly, the early facial processes are hypoplastic from E10.5 in these mice, and subsequently most CNC-derived cranial skeletal elements are impaired or aplastic, including a severely affected mandible associated with aglossia. In addition, deletion of *Shh* from E9.5 in an *Nkx2.5*-expressing cell lineage (*Nkx2.5Cre*; *Shh^flx/ko^*), which overlaps that of *Shh* in oropharyngeal endoderm (Moses et al., 2001) also results in micrognathia and aglossia (Billmyre and Klingensmith, 2015). Whilst these studies demonstrate a clear role for *Shh* in establishment of the tongue primordia, they do not address potential later roles for *Shh* in patterning of the tongue tissues.

In this study, we have utilized a series of genetic approaches to investigate the relevance of Shh signaling during tongue formation, focusing on the source, timing and effect on receiving tissues. We find that production of Shh ligand in the lingual epithelium up to E12.5 is crucial for normal patterning of the intrinsic musculature and that this occurs through signaling to CNC cells to differentiate lingual tendons. Our studies provide the understanding of tongue phenotypes seen in some patients with ciliopathies, such as *orofacial-digital-syndrome-1*, which include tongue clefting, cystic and hamartomatous changes, as well as cleft palate.

## Results

### Shh signaling is required for intrinsic muscle organization and tendon formation in the developing tongue

*Shh* is an early marker for tongue development, being expressed in dorsal epithelium of the lateral lingual swellings from E10.5 and upregulated in lateral regions of the primitive tongue by E11.0 before localizing to early placodal epithelium from E12 (Jung et al., 1999). We have investigated the contribution of *Shh* during tongue development with temporal abrogation of function in mutant mice. Specifically, we crossed *CreER^TM^* mice (Hayashi and McMahon, 2002) with a line harboring a conditional (floxed) *Shh^c^* allele with *loxP* sites flanking exon 2 of the endogenous *Shh* locus (Dassule et al., 2000). After recombination, approximately half of the N-terminal signaling domain is removed from the Shh protein, rendering it non-functional. Following the maternal administration of tamoxifen between E10.5-12.5 and analysis three days later, there was evidence of disruption in anatomy of the tongue in mutant mice when compared to wild type (WT) (Figure 1A-D). In particular, significant disorganization of the intrinsic musculature within the tongue dorsum was present, which was grossly dependent upon the timing of signal loss (Figure 1E-H). In mice treated with tamoxifen at E10.5, the normal striated architecture of the intrinsic muscles was completely lost, whilst deletion at later stages resulted in a progressively less severe myogenic patterning phenotype (Figure 1E-H). However, *in situ* hybridization for the myoblast marker *Myod1*, in WT and mutant embryos suggested that myoblast differentiation had occurred in both the intrinsic and extrinsic musculature, even in the most severely affected mutants, but that organization of the myotubes was defective (Figure 1I-L).

**Figure 1.**
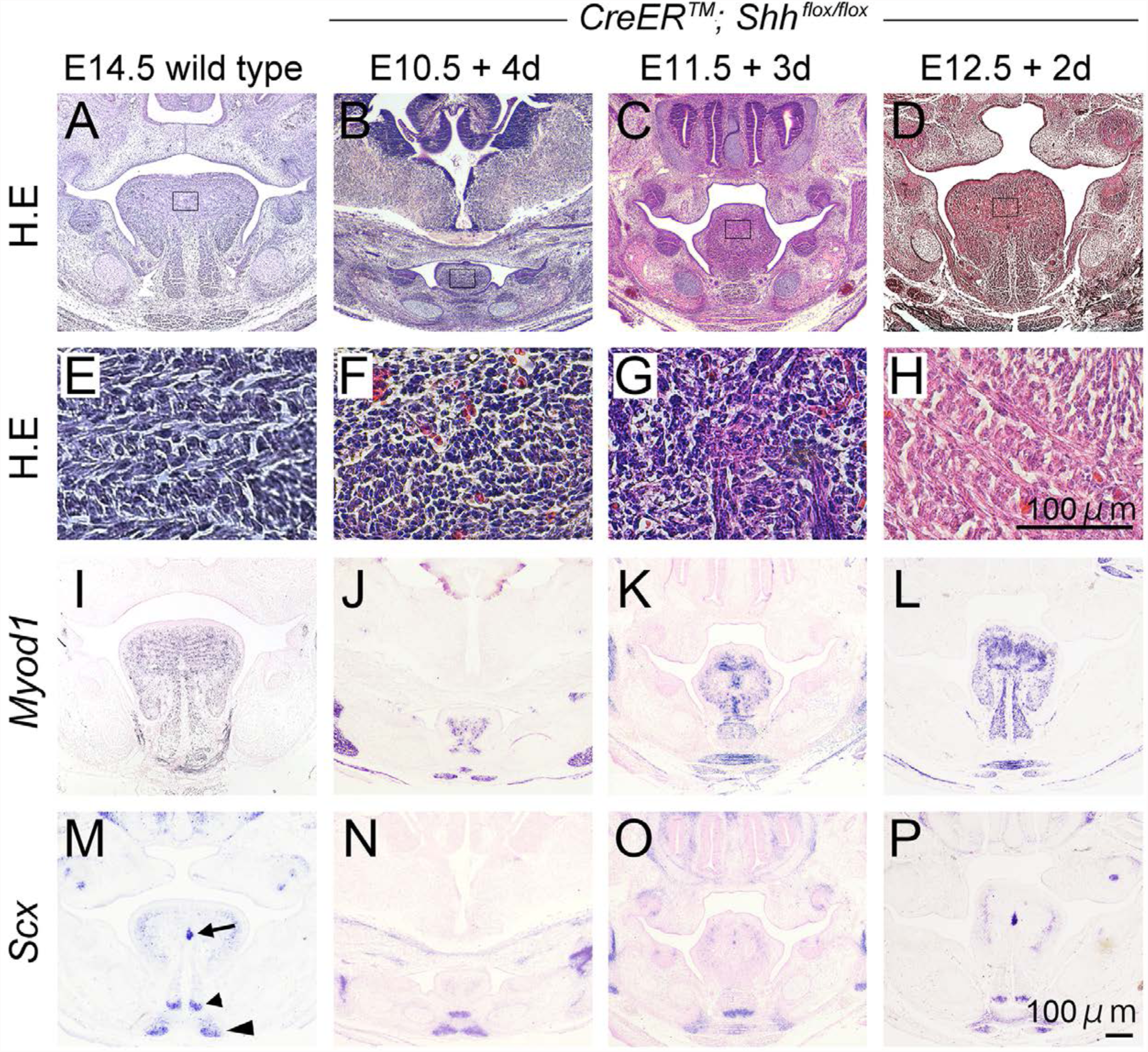
Temporal loss of Shh function produces graded effects on tongue development. (A-D) H.E stained frontal histological analysis of the developing tongue in (A) E14.5 WT and (B-D) *CreER^TM^; Shh^fl/fl^* embryos injected with tamoxifen at E10.5, E11.5, E12.5 and harvested 3 days later, respectively; (E-H) Highlight of intrinsic muscle organization in the tongue dorsum identified within the black rectangles (A-D); (I-P) ISH of *Myod1* (I-L) and *Scx* (M-P) on WT (I, M), and *CreER^TM^; Shh^fl/fl^* embryos that received tamoxifen at E10.5, E11.5, E12.5 and were harvested 3 days later (J, N; K, O; L, P; respectively). *Scx* expression in WT embryos at E14.5 is highlighted in (M) in the lingual septum (arrow), genioglossus (small arrowhead) and geniohyoid (large arrowhead) muscles. Scale bar in P=100μm for A-D and I-P; in H=100μm for E-H.

CNC-derived mesenchyme within the tongue has been suggested to act as a scaffold during the organization of myoblasts and myotubes (Parada et al., 2015). The tongue musculature is patterned through the formation of tendons, which include the aponeurosis and lingual septum, which forms within the tongue dorsum as a flat broad vertical sheath of midline fibrous tissue. We investigated expression of the *Scleraxis* (*Scx*) transcription factor, which is an early marker of tenocyte differentiation (Schweitzer et al., 2001) in WT and mutant embryos at E14.5 (Figure 1M-P). As expected, *Scx* was strongly expressed in the developing midline lingual septum in WT (Han et al., 2012) with more widespread low-level expression in lateral regions of the tongue dorsum. In addition, there was strong bilateral midline expression at sites of tendon formation associated with the paired genioglossus and geniohyoid muscle bellies superiorly and inferiorly, respectively (Figure 1M). In contrast, whilst *Scx* expression was present in the developing genioglossus and geniohyoid tendons of the mutant, there was a loss of expression in the midline septum and significant downregulation in lateral regions of the dorsum in embryos injected at E10.5 (Figure 1N). However, this expression was progressively increased in embryos exposed to later injection at E11.5 and 12.5, respectively (Figure 1O and P). Collectively, these data demonstrate an important role for Shh in mediating normal organization of the intrinsic musculature of the tongue, including tendon architecture.

### Shh in the oropharyngeal epithelium is required for intrinsic muscle organization and tendon formation

The expression of *Shh* in epithelium extending from the oral cavity to the gut is regulated during development by the conserved long-range *cis*-regulatory enhancers, *MRCS1, MFCS4* and *MACS1* located 600 to 900 kb upstream of the mouse *Shh* locus (Sagai et al., 2009). *MFCS4* is responsible for regulating the majority of *Shh* expression in oropharyngeal epithelium from which the incisor tooth bud, pituitary fossa, soft palate, tympanic tube, tongue, epiglottis and arytenoid cartilages have initiated development by E11.0. Targeted deletion of *MFCS4* in mice results in shortening of the soft palate as well as deformation of the epiglottis, arytenoid and tongue, although the size of tongue and mandible are normal (Sagai et al., 2009).

In order to further define the influence of Shh signaling in tongue development we analyzed compound heterozygous *Shh^+/-^; MFCS4^-/+^* mice, where Shh signaling in the developing tongue is clearly decreased by E11.75, as demonstrated by expression of *Shh* and *Ptch1* (Figure 2A-D). At this stage of development, *Myf5*-positive myoblasts have reached the developing tongue dorsum from the occipital somites in WT and *Shh^+/-^; MFCS4^-/+^* embryos (Figure 2F and G), confirming that myoblast migration into the tongue primordium is not affected by loss of Shh signaling. The presence of myoblast differentiation was also confirmed in the WT tongue through detection of *Myod1* in anterior and posterior regions at E11.5 (Figure 2Ha and Hp) with myoblasts present anteriorly expressing *Myod1*, but not desmin at this stage (Figure 2Ha and Ja); whilst those more posteriorly expressed both markers (Figure 2Hp and Jp). Since desmin is a key subunit of the intermediate filaments required for contraction, these observations suggest that lingual myoblast differentiation progresses from posterior to anterior during normal tongue development. This expression pattern was not altered in *Shh^+/-^; MFCS4^-/+^* mutants (Figure 2Ia, Ip, Ka, and Kp). These observations were maintained at E12.5 and confirmed by quantitative RT-PCR (Figure 2E). At E13.5, myotube organization was disrupted and distinct in the superior longitudinal, vertical and transverse intrinsic muscles (Figure 2L and M) compared to lateral regions, where the extrinsic styloglossus muscle runs. In contrast, vascularization and innervation did not appear to be affected in the mutant (Figure 1 - figure supplement 1). As we found in *CreER^TM^*; *Shh^fl/fl^* embryos, *Shh^+/-^; MFCS4^-/+^* mice failed to develop *Scx*-positive tendons for the intrinsic muscles at E13.5 (Figure 2N and O) despite clear expression in the developing genioglossus muscle within the symphyseal region of the mandible (Figure 2N and O arrowheads). *Scx* expression in lateral regions was identified in some *Shh^+/-^; MFCS4^-/+^* embryos (Figure 2O arrows).

**Figure 2.**
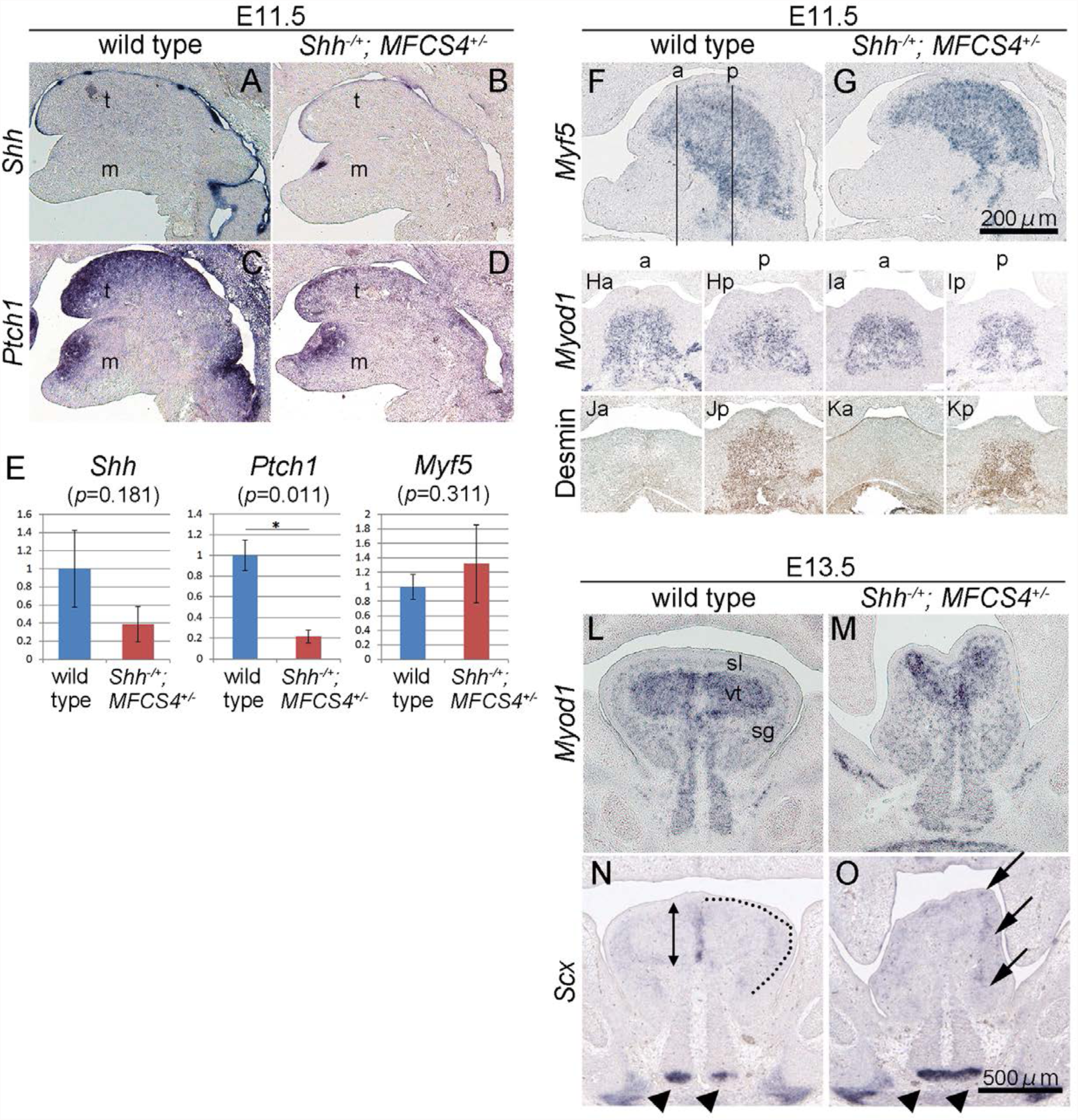
Reduced Shh signaling in the lingual epithelium results in impaired myotube arrangement and tendon formation. (A-D) ISH for *Shh* (A, B) and *Ptch1* (C, D) on sagittal sections of WT littermate (A, C) and *Shh^+/-^; MFCS4^-/+^* (B, D) heads at E11.5. (E) Semi-quantitative RT-PCR analysis of *Shh*, *Ptch1*, and *Myf5* transcription in the tongue of the *Shh^+/-^; MFCS4^-/+^* and the littermate WT at E12.5. (F, G) Myoblast immigration was analyzed by ISH of *Myf5* (F, G) on sagittal sections of E11.5 WT (F) and *Shh^+/-^; MFCS4^-/+^* (G) head. The anterior is to the left. (H-K) Myoblast differentiation was investigated by ISH of *Myod1* (H, I) and IHC of desmin (J, K) at anterior (Ha, Ia, Ja, Ka) and posterior (Hp, Ip, Jp, Kp) region on coronal sections of E11.5 WT (H, J) and *Shh^+/-^; MFCS4^-/+^* (G). The level of the section obtained was indicated as a for the anterior and p for the posterior in F. (L-O) ISH of *Myod1* and *Scx* on coronal section of E13.5 WT (L, N) and *Shh^+/-^; MFCS4^-/+^* (M, O). Arrowheads indicate the short tendon of genioglossus origin at the superior mental spine. The line of the future aponeurosis is drawn as the dotted line in N, which is not consecutive in the *Shh^+/-^; MFCS4^-/+^* (arrows in O). Lingual septum in the litte rmate WT was indicated double arrows in N. m; mandible, t; tongue, sl; superior longitudinal muscle, vt; vertical and transverse muscle, sg; styloglossus. Scale bar for A-K is in G. Scale bar for L-O is in O.

### Shh signaling in the developing tongue targets CNC cells via the primary cilium

Given that myoblast migration and differentiation was not seemingly affected in the absence of Shh signaling in lingual epithelium, but tendon formation specifically was; we hypothesized that the direct target of Shh was CNC cells. We therefore investigated the spatial relationship between *Ptch1*-positive cells, CNC cells and myoblasts in the developing tongue (Figure 3A-E). The detection of beta-galactosidase, *Ptch1* and *Myf5* transcripts in adjacent sections of the developing tongue at E12.5 in *Wnt1-Cre; R26R* embryos indicated that CNC-derived mesenchymal cells expressed *Ptch1* and was therefore a primary target tissue of Shh signal transduction during tongue development.

**Figure 3.**
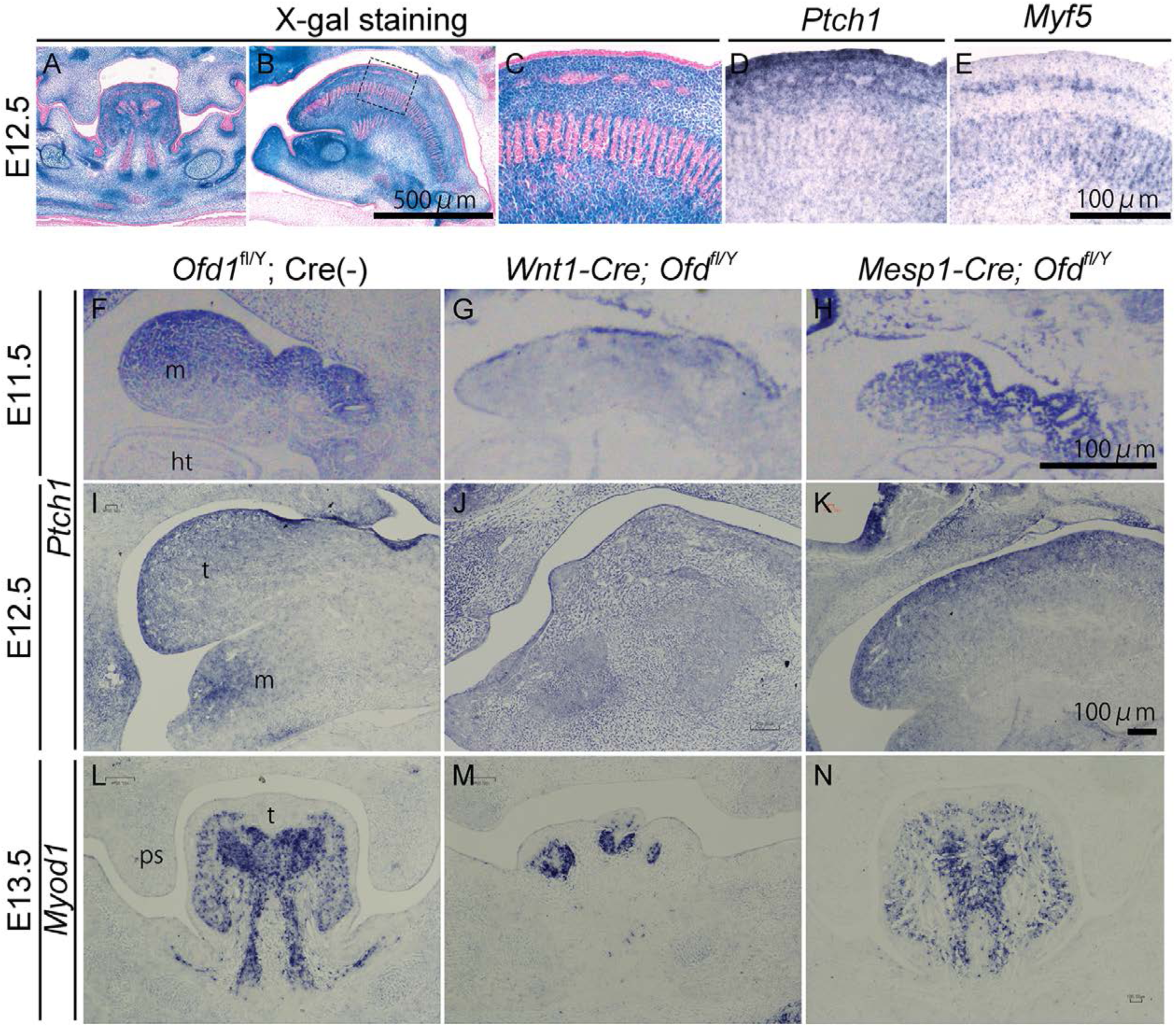
Shh signaling is received by CNC cells for tendon formation and myotube arrangement. (A-C) Lingual CNC-derived mesenchyme was visualized by X-gal staining on frontal (A) and sagittal (B, C) sections of *Wnt1cre; R26R* at E12.5. C is a magnified view of the boxed area in B. (D, E) Shh signal receiving cells were determined by ISH of *Ptch1* (D) and *Myf5* (E) on adjacent sections to B and corresponding region to C. Arrows indicate superior longitudinal muscles in C, D, E. (F-K) The levels of Shh signal is studied by *Ptch1* ISH on sagittal sections of E11.5 (F-H) and E12.5 (I-K) WT (F, I), *Wnt1-Cre; Ofd^fl/Y^* (G, J) and *Mesp1-Cre; Ofd^fl/Y^* (H, K). (L-N) Lingual myotube arrangement is examined by *Myod1* ISH on coronal sections of E13.5 WT (L), *Wnt1-Cre; Ofd^fl/Y^* (M) and *Mesp1-Cre; Ofd^fl/Y^* (N). m; is mandibular process, t; tongue, ht; heart, ps; palatal shelf.

Shh ligand binds to the primary receptor Ptch1, which is located on the primary cilium of receiving cells. Cilia are recognized as key cellular organelles necessary for normal Hh signal transduction in vertebrates, and mutations in the basal body gene *Ofd1* localized on X chromosome lead to a ciliogenesis defect and loss of Shh signal reception (Al-Lami et al., 2016; Ferrante et al., 2006). In human subjects affected by loss-of-function *OFD1* mutations there are multiple craniofacial anomalies, including cleft palate and significant tongue defects, which include clefting, cystic formation and hamartoma. The generation of *Wnt1-Cre; Ofd^fl/Y^* mice with deletion of *Ofd1* in CNC-derived mesenchymal cells led to a reduction of Shh signaling in the tongue, as demonstrated by decreased levels of *Ptch1* expression at E11.5 and 12.5 (Figure 3F and G; I and J, respectively). In contrast, deletion of *Ofd1* in the mesoderm-derived cells (*Mesp1-Cre; Ofd^fl/Y^*) did not affect *Ptch1* expression in the tongue (Figure 3H and K). Significantly, in *Wnt1-Cre; Ofd^fl/Y^* mice there was a complete loss of normal myotube arrangement (Figure 3L and M) as well as severe hypoplastic tongue formation and only clumps of *Myod1*-positive myotubes; whilst in *Mesp1-Cre; Ofd^fl/Y^* mutants, myotube arrangement was largely unaffected (Figure 3N). These observations suggest that lingual tendon formation for intrinsic muscles of the tongue requires Shh signaling from the ventral region of the pharyngeal arches.

### Lingual CNC-derived mesenchymal cell differentiation but not proliferation is affected in *Shh^+/-^; MFCS4^-/+^* mice

We further analyzed CNC cell differentiation in *Shh^+/-^; MFCS4^-/+^* mice. *Sox9* is a marker for CNC cells as well as a common representative transcription factor for chondrocyte, ligament cell and tenocyte differentiation (Spokony et al., 2002; Mori-Akiyama et al., 2003). Upon Bmp-ligand stimulation, mesenchymal cells differentiate into either cartilaginous or tendinous fates by maintaining or decreasing *Sox9* transcription, respectively (Sugimoto et al., 2013. In WT tongues, there was an increase in *Sox9* transcription between E11.75-12.75, with strong expression in the future lingual septum-forming region, that has decreased by E13.75 (Figure 4A, C, and E). In the mutant, *Sox9* was only weakly expressed at E12.75 (Figure 4B, D, and F). Low levels of *Scx* expression was detected from E11.75 in the WT tongue in posterior regions, but not anteriorly (Figure 4G and data not shown). At E12.75, an upregulated M-shaped pattern of expression was seen (Figure 4I), which coincided with the expression pattern in whole mount (Figure 4K). In contrast, there was little evidence of *Scx* transcripts at E11.75 in *Shh^+/-^; MFCS4^-/+^* embryos (Figure 4H) and only faint expression at E12.75 (Figure 4J). Whole mount *in situ* hybridization of *Scx* in E12.75 *Shh^+/-^; MFCS4^-/+^* embryos was consistent with these observations (Figure 4L). Collectively, these observations suggested that during tenocyte differentiation from CNC-derived lingual mesenchymal cells there is a transition stage at which, they express both *Sox9* and *Scx*.

**Figure 4.**
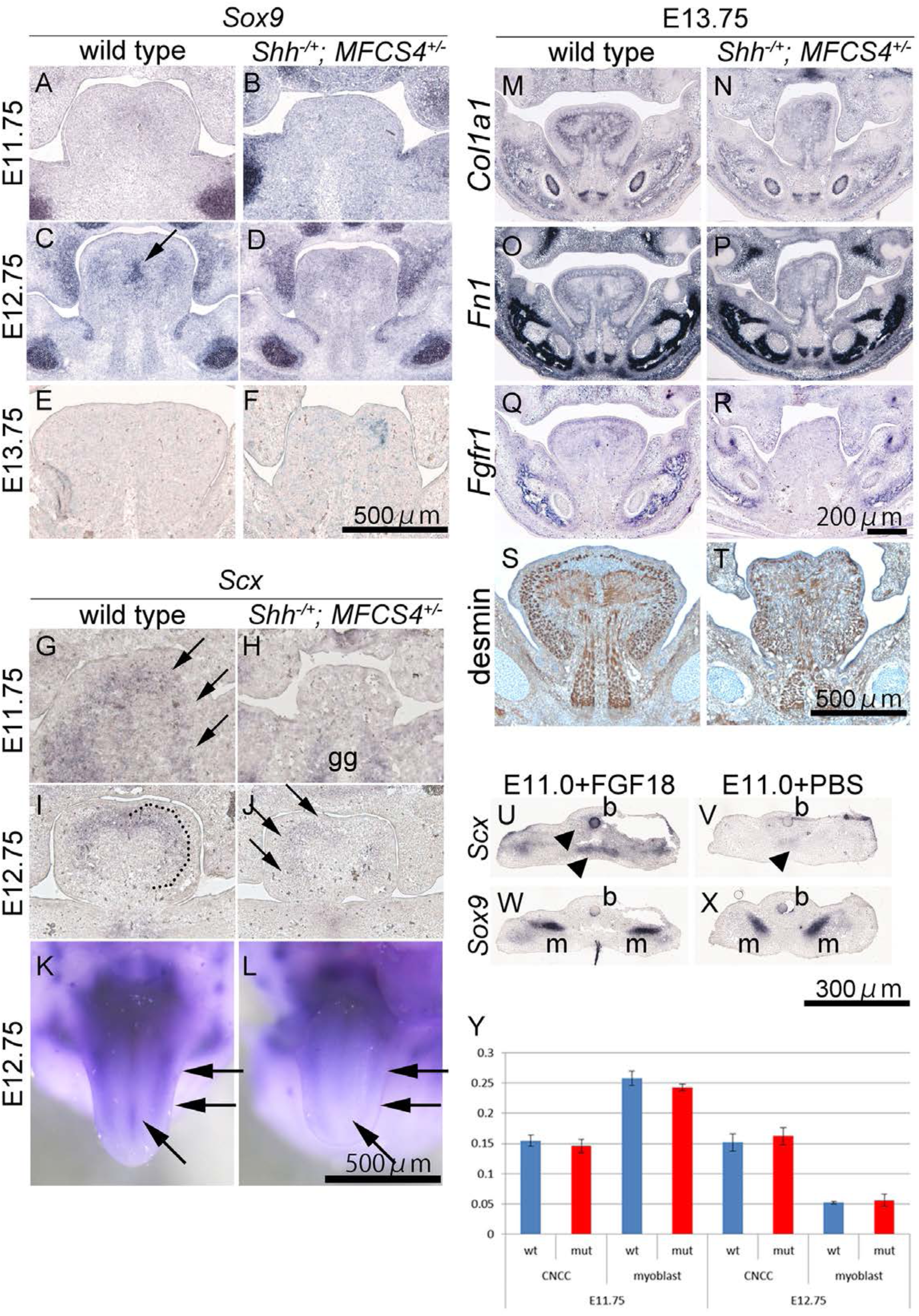
Shh signaling is responsible for differentiation of lingual CNC cells. (A-F) Detection of a mesenchymal marker *Sox9* transcripts on coronal sections at E11.75 (A, B), 12.75 (C, D) and 13.75 (E, F) in WT (A, C, E) and the *Shh^+/-^; MFCS4^-/+^* (B, D, F). *Sox9* expression in the future lingual septum is indicated by arrow in C. (G-L) Stage specific expression of *Scx* was detected on coronal sections of the tongue by ISH (G-J) and whole mount ISH (K, L) at E11.75 (G, H) and E12.75 (I-L) in WT (G, I, K) and *Shh^+/-^; MFCS4^-/+^* (H, J, L). The expression in the future aponeurosis and the lingual septum is indicated by arrows and the dotted line. (M-R) Lingual mesenchyme differentiation was investigated by ISH of *Col1a1* (M, N), *Fn1* (O, P), and *Fgfr1* (Q, R) and IHC of desmin (S, T) on coronal sections of the WT (M, O, Q, S) and *Shh^+/-^; MFCS4^-/+^* (N, P, R, T). (Y) Lingual CNC-derived and mesoderm-derived mesenchymal cell proliferation index was analyzed at E11.75 and E12.75 in WT and *Shh^+/-^; MFCS4^-/+^*. gg; genioglossus muscle. (U-X) E11.0 Mandible after 24h organ culture with FGF18- or PBS-soaked beads were subjected to ISH for *Sox9* and *Scx*. Ectopic expression in the mesenchyme and enhanced expression of *Scx* in genioglossus muscle (arrowheads) were found in the tongue with FGF18 (U). Sox9 expression was not altered by FGF ligand application (W, X). b; beads, m; Meckel’s cartilage.

In accordance with the *Scx* expression pattern, *fibronectin* (*Fn1*) and *type I collagen* (*Col1a1*), and *fibroblast growth factor receptor 1* (*Fgfr1*) were transcribed clearly in the WT aponeurosis and septum (Figure 4M, O, and Q, respectively) whereas expression in the *Shh^+/-^; MFCS4^-/+^* tongue was weak and not well patterned (Figure 4N, P, and R). *Desmin* expression was observed in both WT and *Shh^+/-^; MFCS4^-/+^* tongue at E13.75 although it was disorganized in the latter (Figure 4S and T), which suggested relatively normal myotube formation. Being suggested from the previous reports (Brent et al., 2003, Brent et al., 2005, Sukegawa et al., 2000, Du et al., 2016), we hypothesized Fgf signaling also contribute to tenocyte differentiation. To address this issue, we applied recombinant human FGF18 in the developing tongue at E11.0 and run organ culture. FGF18 induced stronger expression of *Scx* in genioglossus, as well as the strong broad expression in the tongue mesenchyme (Figure 4U and V). *Sox9* transcription was not altered by FGF18 application (Figure 4W and X).

We also studied cell proliferation activity in the early tongue mesenchyme at E11.75 and 12.75 in WT and *Shh^+/-^; MFCS4^-/+^* embryos during which, tenocyte specification occurs in CNC cells (Figure 4Y). Proliferative activity of CNC-derived mesenchyme did not change between E11.75 and E12.75, whereas that of myoblasts was significantly decreased in both WT and *Shh^+/-^; MFCS4^-/+^* embryos, suggestive of active myotube formation during this period. In contrast, no differences in cell death were found in WT and *Shh^+/-^; MFCS4^-/+^* embryos between E11.75 and 13.75 (data not shown).

### A functional tongue is required for secondary palate formation

Previous studies have shown that approximately 14% of *MFCS4^-/-^* mice exhibit cleft palate (Sagai et al., 2009), whilst compound heterozygous *Shh^+/-^; MFCS^+/-^* mice have full penetrance cleft palate and a short soft palate (Figure 5 - figure supplement 2). Histological observation during palatogenesis revealed that palatal shelf elevation was disturbed in the compound *Shh^+/-^; MFCS^+/-^* heterozygote (Figure 5A-F). It has been suggested that palatal shelf elevation and orientation requires coordination between intrinsic factors within the shelves themselves and extrinsic factors such as the descent of the tongue due to embryonic muscle movement, which would provide space in the oral cavity for palate rotation (Iseki et al., 2007). In *Shh^+/-^; MFCS4^+/-^* mice, the palatal shelves were able to elevate and fuse in the midline (n=3) when explanted for organ culture at E13.5, confirming that the cleft palate phenotype resulted from hindrance by the tongue (Figure 5G and H).

**Figure 5.**
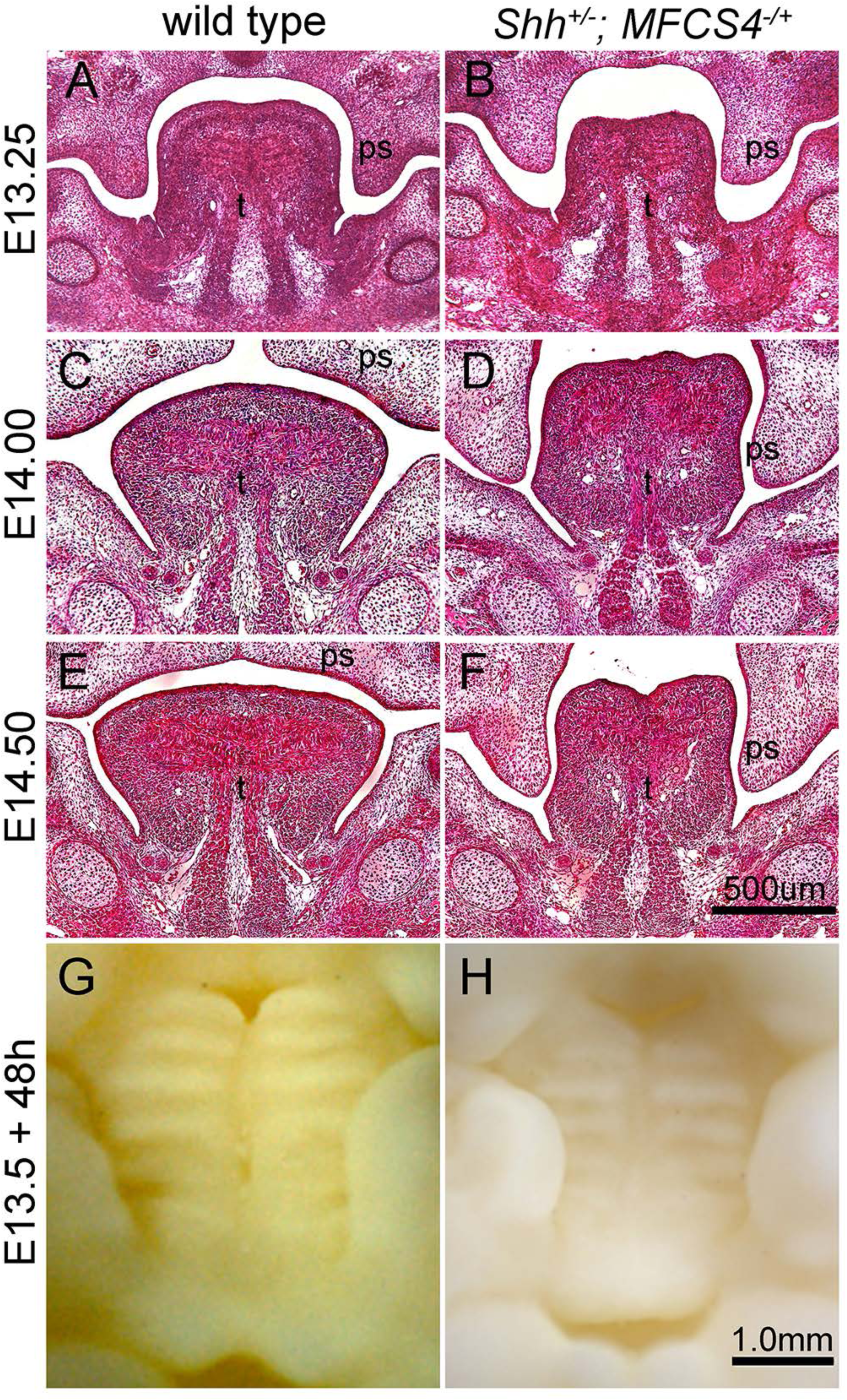
Abnormal tongue formation induces cleft palate. (A-F) H.E staining on frontal sections of WT (A, C, E) and *Shh^+/-^; MFCS4^-/+^* (B, D, F) embryos at E13.25 (A, B), E14.0 (C, D) and E14.5 (E, F). (G, H) Aboral view of the secondary palate after organ culture of E13.5 maxilla of WT (G) and *Shh^+/-^; MFCS4^-/+^*(H). ps; palatal shelf, t; tongue

Micro computed tomography (μCT) analysis on neonatal skeletons of compound heterozygous *Shh^+/-^; MFCS4^-/+^* mice showed a variety of skeletal defects in the maxillary region (Figure 5 - figure supplement 3). In severe cases, an unfused sphenoid, missing premaxilla and vomer and hypoplastic sphenoid tympanic bones were found. However, no defects were identified in the mandible.

## Discussion

A number of signaling pathways have been implicated in tongue formation but the timing and tissue interactions controlling muscle formation have not been well established (reviewed in Parada et al., 2012). *Shh* is expressed in the oropharyngeal epithelium and disruption of signaling in the pharyngeal endoderm prior to tongue morphogenesis (Billmyre and Klingensmith, 2015) or a loss of Hh responsiveness in CNC cells prior to migration (*Wnt1-Cre; Smo^flox/KO^*) (Jeong et al., 2004) both result in aglossia. However, the phenotypic severity associated with both these mouse models precludes analysis of later patterning events in tongue development.

In this study, we show that Shh signaling directly induces formation of the lingual septum derived from CNC cells and that regulation through the *MFCS4* enhancer activated by E11.0 is essential for this process to occur. Removal of Shh signaling throughout the embryo from E10.5 by CreER-mediated site-specific recombination resulted in impaired myotube arrangement within the intrinsic musculature, which was recapitulated in *Shh^+/-^; MFCS4^-/+^* embryos. These results suggest that Shh from the oropharyngeal epithelium induces the tongue primordium before E10.5 and that subsequent signaling is required for differentiation of the lingual septum during internal lingual muscle patterning (Figure 6).

**Figure 6.**
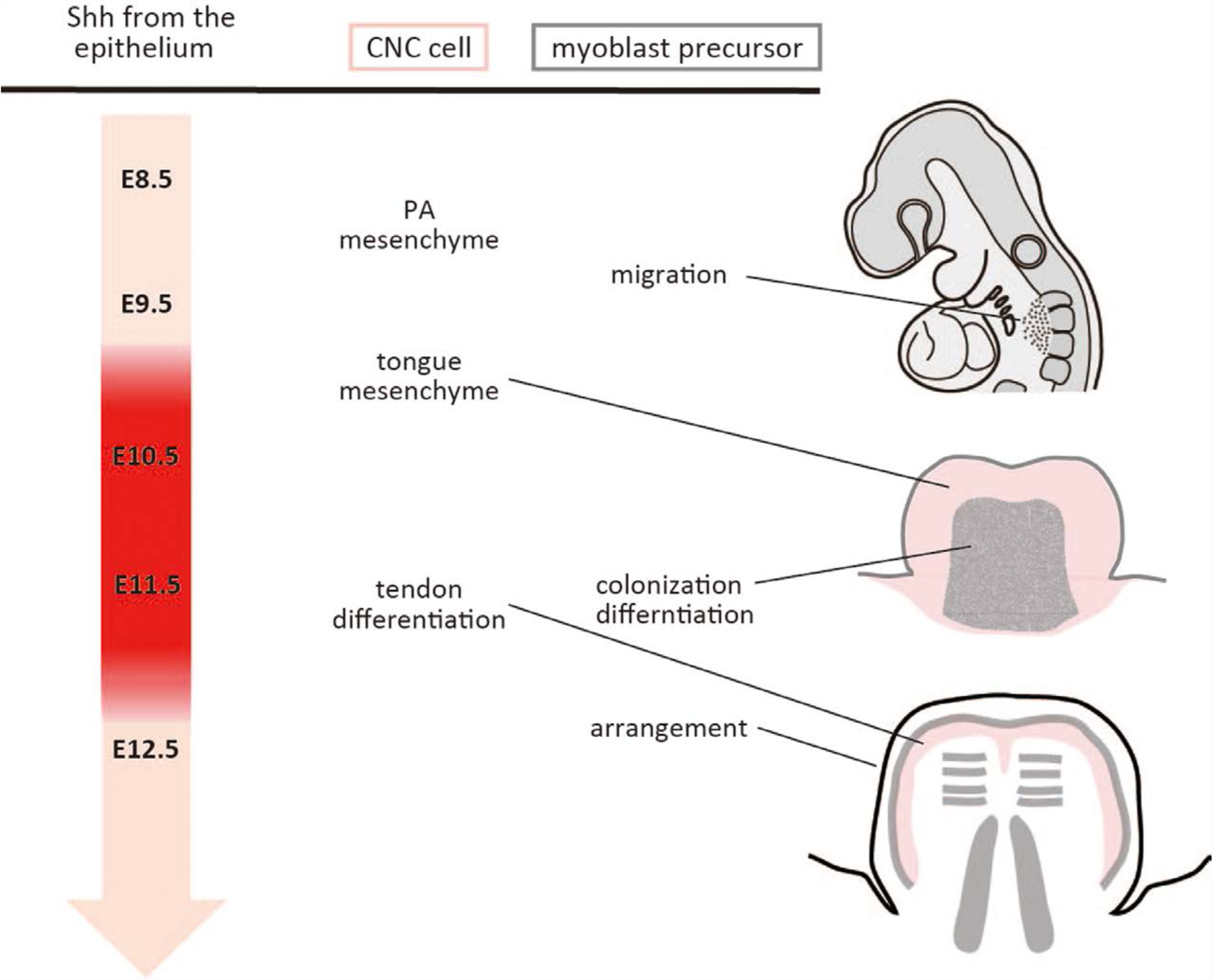
Schematic representation of Shh signaling from the oropharyngeal epithelium during tongue development. The embryonic stage colored in red requires Shh synthesized in the oropharyngeal epithelium. Shh influences CNC-derived mesenchyme (pink) to initiate tongue development and lingual tendon formation during the arrangement of intrinsic lingual muscles.

Our observations from both *CreER^TM^;* Shh*^flox/flox^* and *Shh^+/-^; MFCS4^-/+^* mice suggest that myoblast colonization and differentiation are not affected by altered Shh signaling after the initiation of tongue formation, which was further supported by the tongue phenotype associated with *Mesp1-Cre;Ofd^fl/Y^* mice. We found no difference in myogenic cell lineage proliferation patterns between WT and *Shh^+/-^; MFCS4^-/+^* mice, although Shh signaling is often associated with the regulation of cell proliferation (Ulloa and Briscoe, 2007; Xavier et al., 2016; Ertao et al., 2016; Coni et al., 2017). It appears that Shh secreted from the lingual epithelium after the initiation of tongue formation mostly functions to influence tenocyte differentiation within the internal lingual musculature. Although previous studies have suggested that CNC cells are responsible for the initiation of tongue formation through transduction of the Shh signal (Jeong et al., 2004), it is not clear if myoblasts play a role in this event or they are affected by impaired Shh signaling. *Pax3*-positive muscle progenitors initiate migration around E9.75 and reach the pharyngeal region by E11.5 (Relaix et al., 2004). *Wnt1-Cre; Smo^flox/-^* mice show a cluster of apoptotic cells at the mandibular midline at E9.5, which consequently leads to aglossia. Therefore, it is intriguing to reveal the association between migration of lingual myogenic precursors into the pharyngeal arches and the early stages of tongue formation. There is evidence from studies of TGF-ß signaling during murine tongue formation of a complex interplay between CNC and myogenic precursor cells during this process (Hosokawa et al., 2010; Han et al., 2012; Iwata et al., 2013; Song et al., 2013).

Removal of *Ofd1* in CNC cells (*Wnt1-Cre; Ofd^fl/Y^*) resulted in a hypoplastic tongue and a complete disruption of myotube arrangement; whereas loss of Smo in CNC cells leads to aglossia (Jeong et al., 2004). The *Ofd1* gene is located on the X-chromosome and encodes a component of the centrosome and basal body of primary cilia, a key mediator of Hedgehog signaling (Satir et al., 2010). Smo is a Hedgehog signal mediator, which facilitates the release of Gli transcription factors from the Hedgehog repressor Suppressor-of-Fused (SuFu). Impaired function of the cilium will result in a variable effect on signal transduction, depending upon the molecular nature (Bang and Anderson, 2015) when compared to loss of Smo, which is absolutely required for signal transduction. Mutations in human *OFD1* cause Oral-facial-digital syndrome (OFD) type I, which includes a spectrum of craniofacial phenotypes including gingival frenulae, lingual hamartomas, cleft palate and significantly, a cleft and/or lobulated tongue (reviewed in Franco and Thauvin-Robinet, 2016).

Tendons represent connective tissues that assemble musculoskeletal tissues and anchor force-generating muscles to the skeleton, which leads to optimal locomotion and mobility in vertebrates. The tendon is required to integrate with muscle and skeleton at the myotendinous junction and enthesis, respectively. The mechanism of tendon formation and attachment has been investigated in the trunk, limb and head (Schweizer et al., 2010; Subramanian et al., 2015). The external tongue muscles originate predominantly from hard tissue outside of the tongue and insert into the tongue dorsum and lingual septum. We have detected diminished expression of Fgfr1 in the lamina propria, future aponeurosis, and lingual septum in the tongue of *Shh^+/-^; MFCS4^-/+^* (Figure 4Q, R). There is evidence that *Scx* expression in the early tongue is dependent upon TGF-β signaling from CNC cells but not myogenic precursors (Han et al., 2012; Han et al., 2014). The contribution of Shh signaling in tendon formation has been reported in a variety of developmental systems but not the tongue. In chick axis tendon formation, *Shh* expressed from ventral midline structures such as the floor plate and notochord inhibits induction of syndetome which is comprised of tendon progenitor cells. However, Shh indirectly induces *Scx* expression through activation of fibroblast growth factor (FGF) expression in the dermomyotome, which promotes *Scx* transcription in the somite (Brent et al., 2003). In support of this, FGF signaling is required for differentiation of tenocyte precursors in mice (Brent et al., 2005). In the chick digestive system, expression of *Scx* in two tendon domains that develop in close relation to the two visceral smooth muscles also depends upon FGF signaling (Guen et al., 2009) and *Shh* expressed from the endoderm is responsible for inhibition of smooth muscle cell differentiation from the common undifferentiated mesodermal-derived mesenchyme (Sukegawa et al., 2000).A recent study has reported that expression levels of some FGFs are increased during lingual septum formation (Du et al., 2016).Our *ex vivo* organ culture of the tongue with FGF ligand has resulted in successful induction of *Scx*.

This concludes that Shh signaling influences tenocyte differentiation through Fgf signaling, but *Sox9* expression without. The interaction between Shh and FGF signaling in tendon formation will require further elucidation to fully understand the molecular mechanisms of tendon development in detail.

The obstruction of palatal closure by the tongue is widely thought to be a major cause of cleft palate in human populations. One possible cause of this is abnormal muscle formation and function in the tongue. There are suggestive cases reported previously (Iseki et al., 2007; Okano et al., 2012; Song et al., 2013) and the present study also suggests this possibility. It is also speculated that micrognathia results in abnormal placement or development of the tongue, which can lead to cleft palate (reviewed in Price et al., 2016). There is currently no consensus amongst researchers regarding the involvement of the tongue and mandible during secondary palate formation. More evidence will be required to further define this complex relationship amongst these diverse structures during formation of the oro-facial region.

## Materials and methods

### Animals

All animal experiments were performed in accordance with protocols certified by the Institutional Animal Care and Use Committee of King’s College London and Tokyo Medical and Dental University. *MFCS4^+/-^* (Sagai et al., 2009), *Shh^-/+^* (Amano et al., 2009), *CreER^TM^* (Hayashi and McMahon, 2002), *Shh^flox/+^* (Dassule et al., 2000), *Ofd1* (Ferrante et al., 2006), *Wnt1-cre* (Danielian et al., 1997), *Mesp1-cre* (Saga et al., 1999) mice were maintained in a C57BL/6N background. *CreER^TM^; Shh^flox/+^* mice were mated with *Shh^flox/+^* mice and pregnant mice received tamoxifen by intraperitoneal injection (75 mg/kg, equivalent to 3 mg per 40g body weight) through the maternal body.

### Histological analyses

Specimen were fixed with Bouin’s solution for hematoxylin-eosin (H.E.) staining or 4% paraformaldehyde (PFA)/PBS, embedded in paraffin (H.E.) or O.C.T. compound (Sakura Finetek, Tokyo, JP for other histological analyses), and prepared to 5 μm-(paraffin) or 12 μm (frozen)-thick sections. For immunohistochemistry (IHC), anti-desmin antibody (clone D33, 413651, Nichirei Biosciences, Tokyo, JP) at x1 dilution, anti-SMA antibody (clone 1A4, A2547-100UL, Sigma, Saint Louis, MO) at x1,000 dilution, anti-BrdU antibody (clone BMC9318, 11 170 376 001, Roche Diagnostics, Basel, CH) at x100 dilution, anti-Myf5 antibody (polyclonal, SAB4501943, Sigma, Saint Louis, MO) at x100 dilution, anti-CD31 antibody (polyclonal, ab28364, Abcam, Cambridge, UK) at x100 dilution, and anti-synaptophysin antibody (clone SY38, ab8049, Abcam, Cambridge, UK) at x100 dilution were used. For visualization, corresponding secondary antibodies, VECTASTAIN ABC Kit (AK-5000, Vectastain, Burlingame, CA) and diaminobenzidine (DAB) were applied, or corresponding fluorescent-conjugated secondary antibody was applied. Hematoxylin and Hoechst 33342 were used for counter staining for DAB and fluorescence, respectively.

For *in situ* hybridization (ISH), specimens were hybridized with digoxygenin-labeled RNA probes specifically designed complementary to the full- or partial- mRNA of *Shh*, *Ptch1, Myf5*, *Sox9*, *Scx, Col1a1, Fn1,* and *Fgfr1* followed by the incubation with anti-digoxygenin-AP conjugate. Nitro blue tetrazolium chloride (NBT) / 5-Bromo-4-chloro-3-indolyl phosphate, toluidine salt (BCIP) were used for colourization. All commercial reagents for ISH were purchased from Roche Diagnostics (Basel, CH). All template DNA for RNA probes were cloned into pTA2 vector (TArget Clone, TAK-101, Toyobo Life Science, Osaka, JP).

For X-gal staining (detection of beta-galactosidase), sections were incubated with 5-bromo-4-chloro-3-indolyl-β-D-galactoside (X-gal) in phosphate buffer (pH7.3) supplemented with 2 mM MgCl_2_, 5mM potassium ferrocyanide (K_4_Fe(CN)_6_-3H_2_O), and 5mM potassium ferricyanide (K_3_Fe(CN)_6_) at 37°C after fixation in 4%PFA. Nuclear Fast Red was used for counter staining.

### Cell proliferation analysis

Sixty minutes before dissection, BrdU at 100mg/ml was injected intraperitoneally to the pregnant mice at 10mg/kg at the designed day. Every 6 of 7 sections through the tongue primordium of a specimen were used for cellular proliferation index study. BrdU incorporation and Myf5 localization were detected by immunohistochemistry. Epithelial cells were determined as external cells than the basement membrane by histological observation. Myoblast cell lineage was determined as Myf5-positive cells, and Myf5-negative cells were assumed to be CNC-derived. Proliferation index was calculated by the number of BrdU-positive cells] / [total number of cells of each population] and statistical significance was examined by *t*-test for more than 3 individual experiments for each genotype.

### Real time RT-PCR

RNA was extracted from the tongue of littermate WT and *Shh^+/-^; MFCS4^-/+^* embryos at E12.5 using Direct-zol RNA MiniPrep Kit (R2050S, Zymo Research, Irvine, CA) following the product protocol. 250ng RNA was transcribed to cDNA by ReverTra Ace (TRT-101; Toyobo Life Science, Osaka, JP). Realtime PCR was performed with LightCycler 480 High Resolution Melting Master (04909631001; Roche, Diagnostics, Basel, CH). Normalized to the *actin, beta* gene and relative expression to the littermate WT was shown. Statistical significance was examined by *t*-test for more than 3 individual experiments for each genotype.

### Organ culture

The dissected maxilla (with palatal shelves) of E13.5 *Shh^+/-^; MFCS4^-/+^* or WT littermates were cultured in the mixture of Dulbecco’s Modified Eagle’s Medium Nutrient Mixture F-12 (Sigma-Aldrich, St. Louis, MO) and BGJb Medium (LifeTechnologies, Grand Island, NY) for 48 hours by supplying 95% O_2_ + 5% CO_2_ at 37°C using rotary culture system.

Heparin-coated acrylic beads (diameter, 125–150 mm; Sigma, St Louis, MO) were soaked in either PBS or in 300 mg/mL recombinant human FGF18 (AF-100-28, PEPROTECH, Rocky Hill, NJ) for at least 1 hour at room temperature. The beads were embedded in the tongue with mandible at E11.0 and the tongue underwent the rotary organ culture system as described above for 24 hours.

### Micro computer tomography images

*Shh^+/-^; MFCS4^-/+^* and their WT littermates at P0 were scanned with X-ray microCT inspeXio SMX-100CT (Shimadzu Corporation, Kyoto, JP) and rendered with VGStudio Max 2.0 (Volume Graphics, Heidelberg, DE). Each figure shown is a representative of more than 3 independent experiments. Pictures paired for comparison were taken and processed under the same condition from the same litter.

## Acknowledgements

This work was supported by JSPS KAKENHI Grant Numbers 19890071, 22592254, 25463130 and 16K11744 to S.O., and 20390510 to S.I., and NIG-JOINT (2008-A, 2009-B7, 2012-B4) to S.I. H.A.A. was funded by Iraq Higher Committee for Educational Development. K.J.L. received the funding from the BBRSC (Grant BB/I021922/1) and MRC (Grant MR/L017237/1)

S.O. and A.B. are co-first, and M.C. and S.I. are co-corresponding authors.

---

**Figure Supplement 1.**
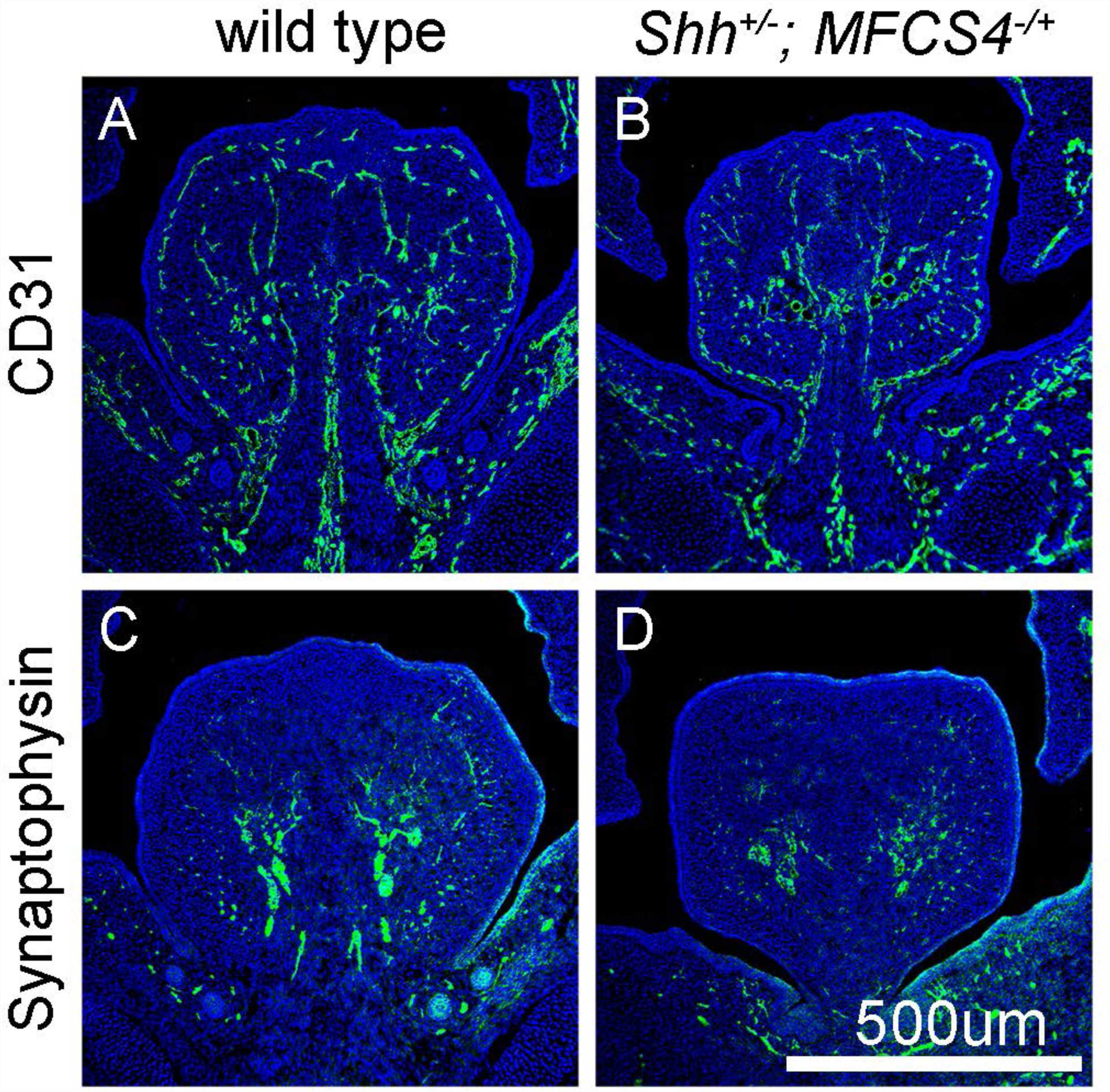
Vascularization and innervation is not affected in *Shh^+/-^; MFCS4^-/+^*. (A, B) Immunofluorescent detection of an endothelial marker CD31 (green) on E13.5 coronal sections of WT (A) and *Shh^+/-^; MFCS4^-/+^* (B). (C, D) Immunofluorescent detection of the neuron marker synaptophysin (green) on E13.5 coronal sections of WT (C) and *Shh^+/-^; MFCS4^-/+^* (D). Nuclei are stained blue by Hoechest. Both vascular endothelial cells (A, B) and pre-synaptic protein (C, D) were detected in a similar manner in WT and *Shh^+/-^; MFCS4^-/+^*.

**Figure Supplement 2.**
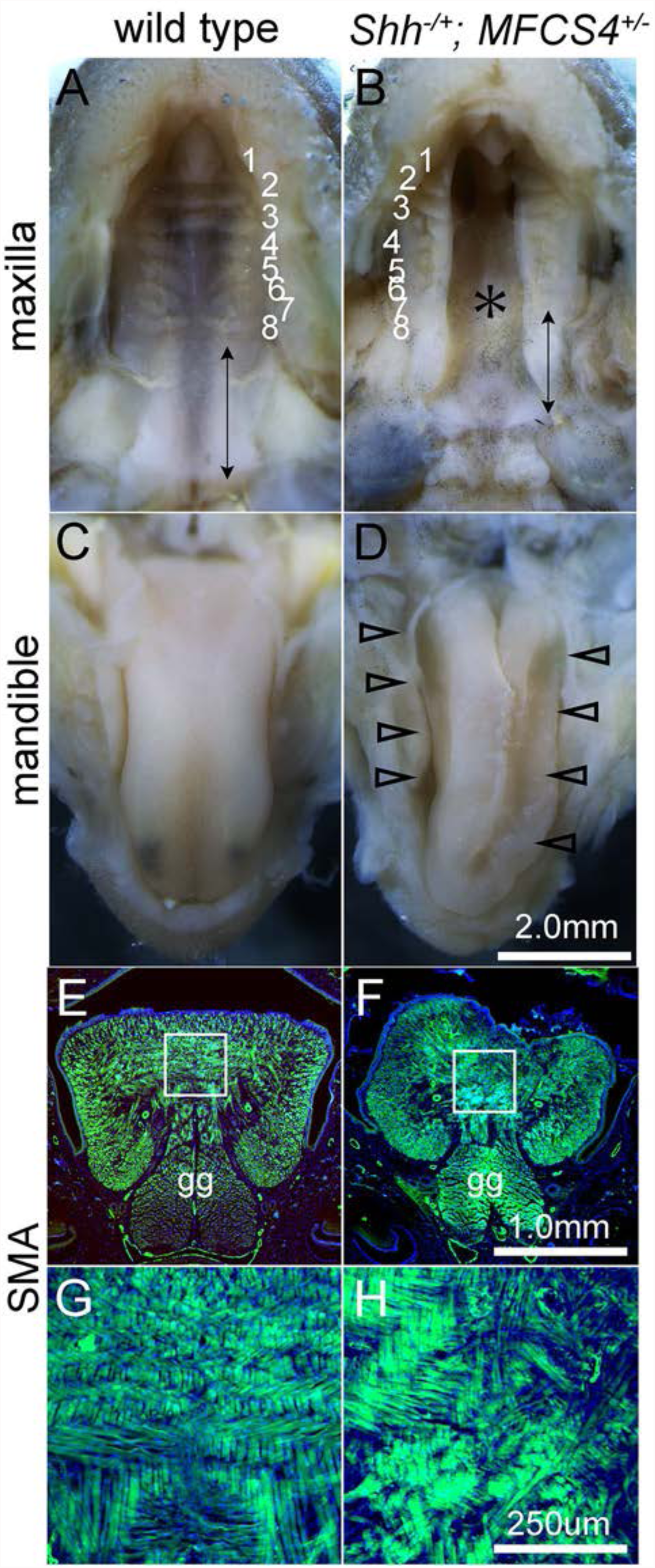
*Shh^+/-^; MFCS4^-/+^* mice exhibit cleft palate and disorganized myotube arrangement at P0. (A-D) Maxillae (A, B) and mandibles (C, D) of WT (A, C) and *Shh^+/-^; MFCS4^-/+^* (C, D). **WT** palate is fully fused in the midline whereas that of *Shh^+/-^; MFCS4^-/+^* is completely cleft (asterisk). Double-headed arrow points the soft palate region and the number is given to each ruga. The *Shh^+/-^; MFCS4^-/+^* also demonstrate short soft palate. The size of the mandible of *Shh^+/-^; MFCS4^-/+^* is similar to that of WT. The tongue has been distorted with grooves that are considered to be the impression by the palatal shelves (arrowheads). *Shh^+/-^; MFCS4^-/+^* exhibit disorganized myotubes arrangement. (E-H) Immunofluorescent detection of a myotube marker smooth muscle actin (SMA) on coronal sections of WT (E) and *Shh^+/-^; MFCS4^-/+^* (F). G and H are magnified views of the boxed area in E and F, respectively. Myotube arrangement in WT was symmetric and both vertical and transverse myotubes are arranged in order. In contrast, *Shh^+/-^; MFCS4^-/+^* myotube arrangement is asymmetric and disturbed, especially in the midline area due to lack of lingual septum formation.

**Figure supplement 3.**
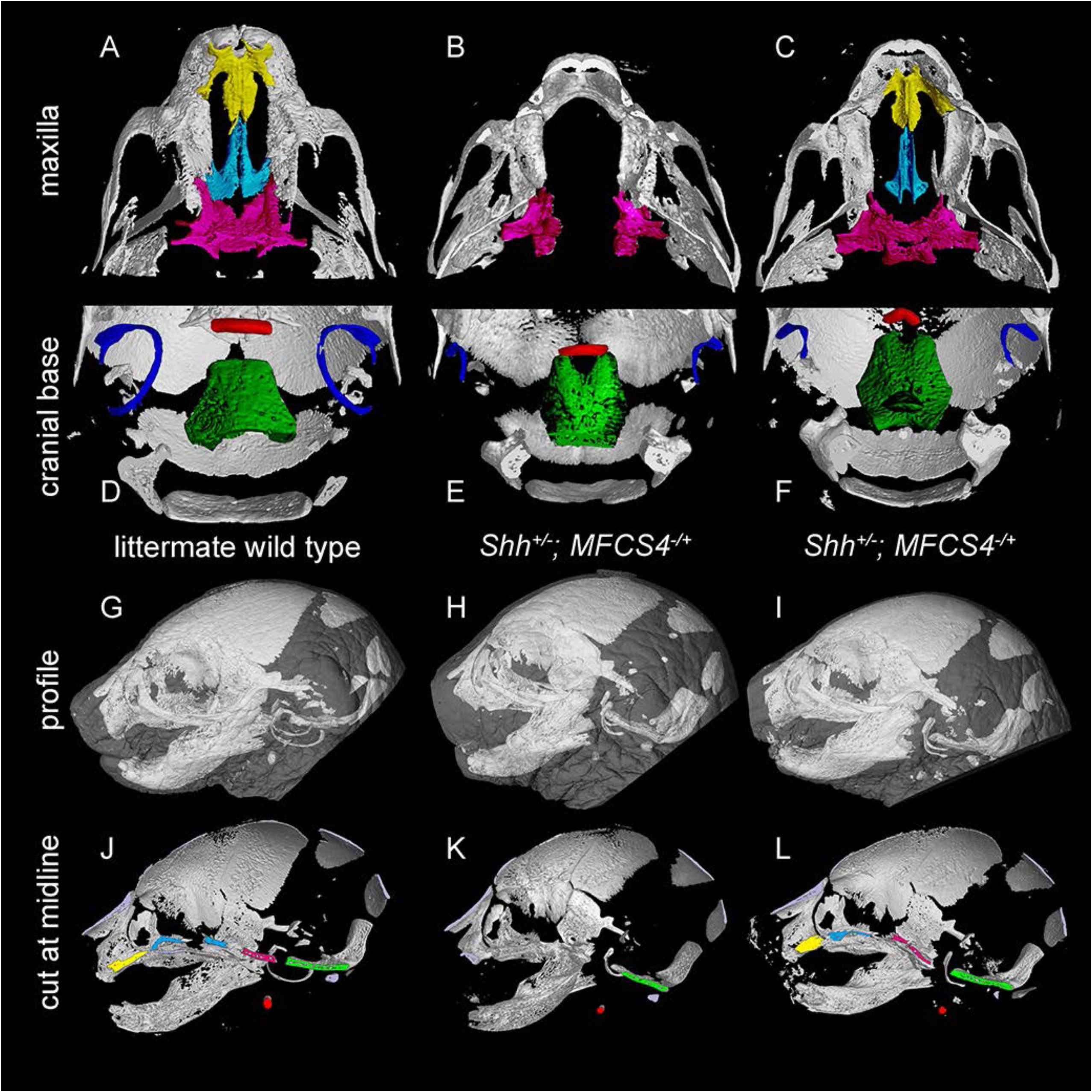
Skeletal defects in *Shh^+/-^; MFCS4^-/+^*. MicroCT scanned maxilla (A-C), posterior cranial base (D-F), profile (G-I), and skeleton cut at midline (H-J) of *Shh^+/-^; MFCS4^-/+^* and the wild type at P0. The *Shh^+/-^; MFCS4^-/+^* with severe phenotype (B, E, H, L) and milder phenotype (C, F, I, K) are shown to be compared with the wild type (A, D, G, J). Smaller or missing prepalatal bone and vomer, premaxilla, basis of sphenoid bone were found in the *Shh^+/-^; MFCS4^-/+^* (A-F, J-L). Hyoid bone and tympanic ring of the *Shh^+/-^; MFCS4^-/+^* were also smaller than those of the wild type. Basoccipital bones of the *Shh^+/-^; MFCS4^-/+^* had a notch at anterior edge where derived from CNC cells (D-F). These phenotypes met with the origin of each bone and active region of MFCS4. Prepalatal bone and vomer; yellow, premaxilla; cyan, sphenoid bone; magenta, basoccipital bone; green, hyoid bone; red, tympanic ring; blue. Lateral view is the merge of skeletal system with soft tissue in grey.

